# VisioWell, an all-in-one high-throughput platform to study haematopoietic cells division

**DOI:** 10.64898/2026.11.21.768298

**Authors:** Tiziana Tocci, Morgane Burq, Eugenio Martinelli, Arianna Mencattini, Leila Perié, Hervé Isambert, Alessandro Donada

## Abstract

Live-cell imaging has the potential to unravel crucial parameters for haematopoietic normal and leukaemic stem and progenitor cells. Yet, this technology requires a combination of technical and analytical skill sets that can be difficult to find in a traditional haematology lab. We present VisioWell, an all-in-one experimental platform specifically designed to measure the early division dynamics of blood stem and progenitor cells, from a simple brightfield long-term acquisition. We combined a simple custom-made microwell array to trap thousands of individual cells with a powerful semi-automatic analytical pipeline, able to segment, correct, and retrieve the division times for the first three sets of divisions. We validated VisioWell with primary cells from human cord blood and with a cellular model of myeloid leukaemia, revealing that CDK4/6 inhibition induces an increase in asynchronous division. By lowering the technical and analytical barriers to live-cell imaging, VisioWell makes label-free, non-destructive analysis of division dynamics accessible to any haematology lab, while preserving cells for downstream fate and functional readouts.

## Introduction

Cell division, together with cell differentiation and cell death, are core parameters of the cellular dynamics of haematopoietic stem and progenitor cells (HSPCs). Their accurate measurements are essential to study normal stem cell differentiation and self-renewal, but also their potential dysregulation in the pathological setting. Live-cell imaging has long been recognized as an appealing non-invasive method for studying cell division of rare and heterogeneous cells like HSPCs [1]. For instance, live-cell imaging revealed that mouse clones retaining stem cell activity took longer to divide [2], and that Stem Cell Factor (SCF) stimulation can change the baseline division kinetics [3]. Moreover, the analysis of live-cell imaging experiments enabled the study of collective cellular dynamics across multiple cells, such as the synchronicity of divisions between sister cells (Fig. 1a). High synchronicity in cell division has been reported for B cells [4], whereas altered correlation patterns have been linked to disease [5–7]. We recently observed that healthy human HSPCs display a strong degree of synchronicity, a property partially disrupted in acute myeloid leukaemia (AML) [8]. These datasets were generated using a low-throughput experimental design (clonal cultures in 96-well tissue vessels) and manually annotated. Limited throughput and time-consuming workflow restrict scientific discovery in HSPC dynamics, particularly the identification of key regulators through screening approaches. Live-cell imaging studies have moved away from manual tracking, and multiple automatic and semi-automatic approaches are now available [9, 10]. They nevertheless require a certain degree of technical expertise, and they are not adapted to label-free datasets, for example, human primary samples. Particularly challenging for label-free datasets are the identification (bounding-box level) [11] and the segmentation (pixel-level) [12] of the relevant events. The Fiji software [13], through the TrackMate plugin [14], enables segmentation via different algorithms, including Cellpose pretrained models [15] and the Trainable WeKa Segmentation algorithms [16]. However, most of those tools are not suitable for high-throughput techniques without additional steps. Another major challenge for live-cell imaging approaches is the motility inherent to blood cells, which makes long-term tracking extremely difficult. Several approaches exist, from physically embedding those cells in methylcellulose [17] to elegant biochemical surface coating that reduces cell motility via specific interactions [18]. We propose Visiowell: an all-in-one solution to perform high-throughput, label-free, single-cell live imaging experiments of non-adherent cells. The platform measures cell division dynamics by automating the extraction of cell division features over hundreds of clonal lineages. VisioWell integrates a microfluidic chip [19, 20] containing thousands of microwells, brightfield time-lapse live-cell imaging, and a segmentation-based pipeline to automatically retrieve the division times (Fig. 1b). We validated VisioWell by analyzing the first three sets of divisions in healthy cord blood HSPCs. We also assessed the impact of different small molecules on AML cell division, highlighting the unexpected impact of CDK4/6 inhibition on division synchrony. These observations pave the way for further studies on the role of cell cycle length in HSPC biology.

**Figure 1:**
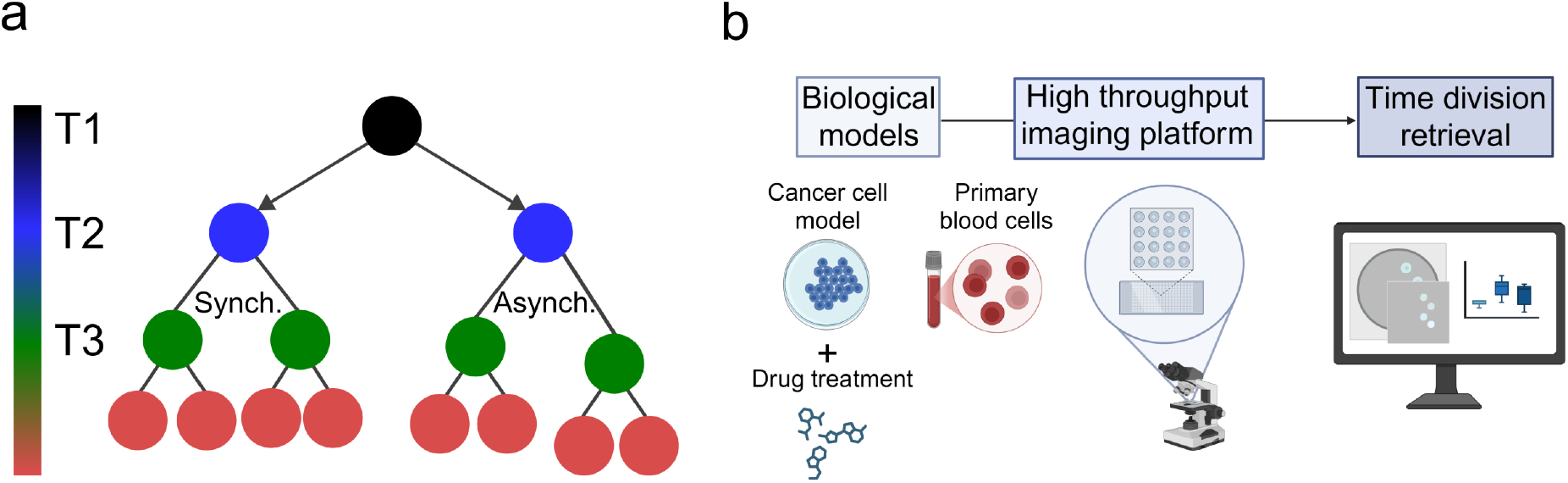
Framework to automatically study cell division dynamics. **a)** Cell family generation tree for the first three divisions. The mother cell is indicated in black, the daughter cells (1st generation) in blue, the cousin cells (2nd generation) in green, and the further generations in red. Since division times are recorded chronologically, only the progenitor identity of the sister cells is known with certainty (indicated by arrows). **b)** VisioWell study design.

## Results

### VisioWell immobilizes individual non-adherent cells, allowing clonal expansion

VisioWell combines a microwell array with an imaging data analysis workflow. This allow to isolates thousands of non-adherent individual cells at the start of the experiment, to univocally identify their descendancy without complex tracking algorithm, or by reducing cell motility [1]. We produced the microwells array by hot-embossing of a biocompatible thermoplastic elastomer named Flexdym [21] (Fig. 2a). We selected a microwell density of 1225 microwells/cm^2^, with each microwell providing sufficient space for approximately 30 cells. The upper arc-shaped design of the microwell promotes rapid cell sedimentation after seeding (1-2 hours), without allowing them to escape from the microwell (Fig. 2b, Fig. S1a).

**Figure 2:**
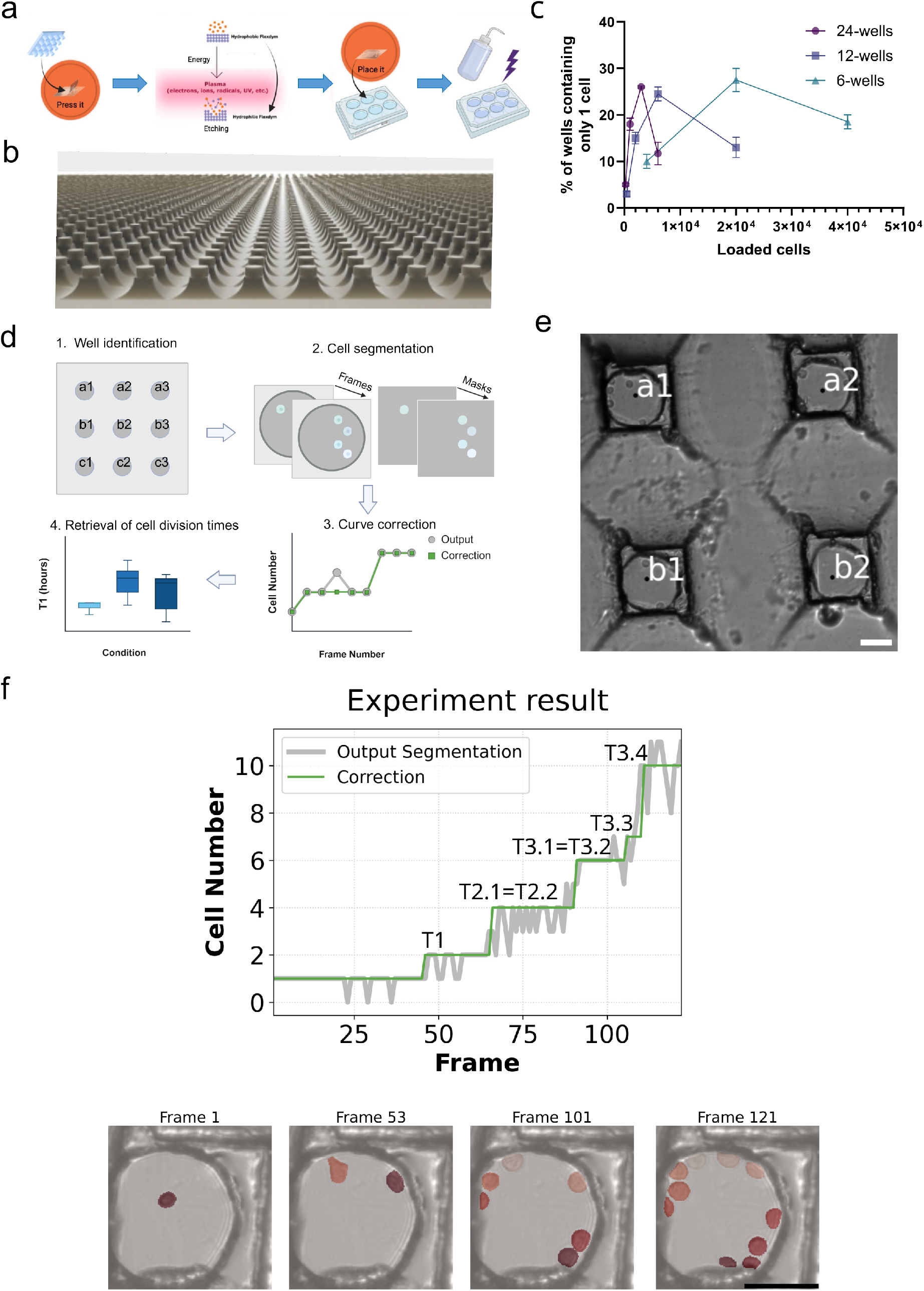
The visioWell platform: microwell array design and computational pipeline. **a)** Schematic of the microarray production workflow. The microwells are printed into a pre-cut flexdym sheet by hot embossing. The microchip is then etched using a plasma cleaner and placed in a culture plate well. The culture plate is then sterilised (with ethanol and/or UV light). **b)** Rendering of the brass mould surface (3D Autodesk Fusion), displaying the micropillar structures for microwell printing. **c)** Cell seeding efficiency, represented as the percentage of wells displaying only one cell, relative to the total number of seeded cells. **d)** Data analysis algorithm workflow. **e)** Example of well identification and naming. Crop on four microwells. Scale bar: 50 *µm*. **f)** Example of cell identification and cell number time series curve correction. The segmentation output is reported in gray. The division times are retrieved from the corrected curve, in green. Distinct segmented cells are indicated with a different shade of red in the frames. Scale bar: 50 *µm*.

The microwell array is compatible with standard 6-, 12-, and 24-well tissue culture plates. Because these plates are compatible with most Fluorescence-Activated Cell Sorting (FACS) instruments, VisioWell can be integrated into existing protocols for rare samples or cell populations that require a FACS sorting step. For each format, we measured the optimal cell concentrations for uniform seeding, to maximise the proportion of microwells containing a single cell (Fig. 2c). The optimal loading efficiency, for the three formats, was between 25% and just below 30%. As an example, for a 12-well single array, containing approximately 3350 microwells, we can analyse up to 837 individual clones in parallel.

### VisioWell imaging process pipeline automatically retrieves cell division times

To enable high-throughput quantification of clonal cell dynamics, we developed an automated analysis pipeline that identifies individual microwells, retrieves cell numbers over time, and uncovers cell division events (Fig. 2d, Fig. S1b). Microwells were automatically detected and indexed according to a grid-based framework (Fig. 2e). Only wells containing a single cell at the start of the experiment were retained to preserve clonality (18974 microwells, from three independent experiments) and we additionally filtered to retain only wells for which at least a division was detected (11915 microwells from three independent experiments).

The pipeline successfully segmented cells across sequential image frames using a finetuned CellposeSAM model [22](Fig. 2f) and generated cell count curves for each microwell (Fig. 2f). To improve robustness, a change-point detection algorithm [23] corrected transient fluctuations in cell-count estimates, yielding stable cell-count curves from which division events could be identified (Fig. 2f). Based on the reconstructed clonal trajectories, the algorithm inferred the timing of individual cell divisions automatically, assuming that all sister cells (1st generation) completed their divisions before the cousin cells (2nd generation) started dividing. This assumption was previously verified [8]. In addition to cell division events, the pipeline identified cell death events through decreases in cell counts, consistent with the confinement of cells within individual microwells (Fig. S1a). These events corresponded to specific morphological changes observed in the frames, including loss of cellular integrity and motility (Fig. S1c). Intermediate outputs could be inspected through a dedicated Streamlit interface (Fig. S1d), and all measurements were consolidated into a unified dataset for downstream quantitative analyses (Fig. S1b).

### VisioWell quality control selects high-quality microwells

To assess the automated image analysis performance, we defined three quality indices, *OscIdx*, *OscIdx*8 and *TrustIdx*. They flag microwells affected by segmentation errors due to poor focus (e.g. uneven surface, air bubbles trapped under the microarray), cell culture debris that are not unequivocally recognized as cells, and cell death. *TrustIdx* takes into account quality checks for the entire acquisition, such as the maximum number of cells at the end of the experiment and the delays between consecutive divisions. *OscIdx* measures the number of oscillations for the whole acquisition, while *OscIdx*8 measures the number of oscillations until the last frame where we can detect a maximum of 8 cells. If the microwell never reaches 8 cells, then *OscIdx* = *OscIdx*8. We assessed the three index distributions across all experiments: we analysed isolated human CD34+ cells from cord blood and leukemic CD34+ cells from OCI-AML8227 cell cultures [8, 24]. After excluding microwells for which no division was detected, we observed that *OscIdx* and *OscIdx*8 distributions were right-skewed, while the *TrustIdx* distribution was left-skewed (Fig. 3a). We then selected a subset of 35 microwells across all datasets, and we manually annotated the divisions to create a ground truth dataset. We then compared those results to the automatic detection output and assessed their correlation. On this subset, *OscIdx* and *OscIdx*8 were highly correlated (Fig. 3b), demonstrating that oscillations measured up to the third division were representative of the quality of the whole experiment and could be used as a robust quality metric. Conversely, *OscIdx*8 and *TrustIdx* were highly anti-correlated (Fig. 3b). To determine the existence of a threshold able to meaningfully separate low-from high-quality microwells, we computed the *MAE*8 (Mean Absolute Error up to the last frame with eight cells), fundamentally comparing the cell numbers from the automatic detection with the manually annotated results (the ground truth). Microwells with *OscIdx*8 *<* 0.2 or *TrustIdx >* 0.8 showed significantly lower *MAE*8 than those above threshold (Fig. 3c), supporting *OscIdx*8 and *TrustIdx* as practical, annotation-free quality filters. To validate the thresholds further, we measured their performance: first, we selected the lower-quality microwells from the 35 (*MAE*8 *>* 0.3, 15 of 35). The indicators respective area under the receiver operating characteristic AUROC (0.81 for *OscIdx*8, 0.79 for *TrustIdx*) and area under the precision-recall curve AUPRC (0.70 for *OscIdx*8, 0.66 for *TrustIdx*) scored positively against a chance level of 0.43 (Fig. S2a). At the chosen thresholds, 80% of unreliable microwells were flagged (12/15), at the cost of discarding 25% (*OscIdx*8) and 30% (*TrustIdx*) of reliable ones, consistent with their conservative purpose. Results were similar for *MAE*8 *>* 0.4 (9 out of 35) with AUROC 0.79 (*OscIdx*8) and 0.77 (*TrustIdx*) and with 89% of unreliable microwells flagged (8/9) at the cost of discarding 35% (*OscIdx*8) and 38% (*TrustIdx*) of reliable ones (Fig. S2b). Cross-validated threshold selection (repeated 5-fold) gave median thresholds of 0.13 (*MAE*8 *>* 0.3) and 0.21 (*MAE*8 *>* 0.4) for *OscIdx*8 and 0.79 (*MAE*8 *>* 0.4) and 0.87 (*MAE*8 *>* 0.3) for *TrustIdx*.

**Figure 3:**
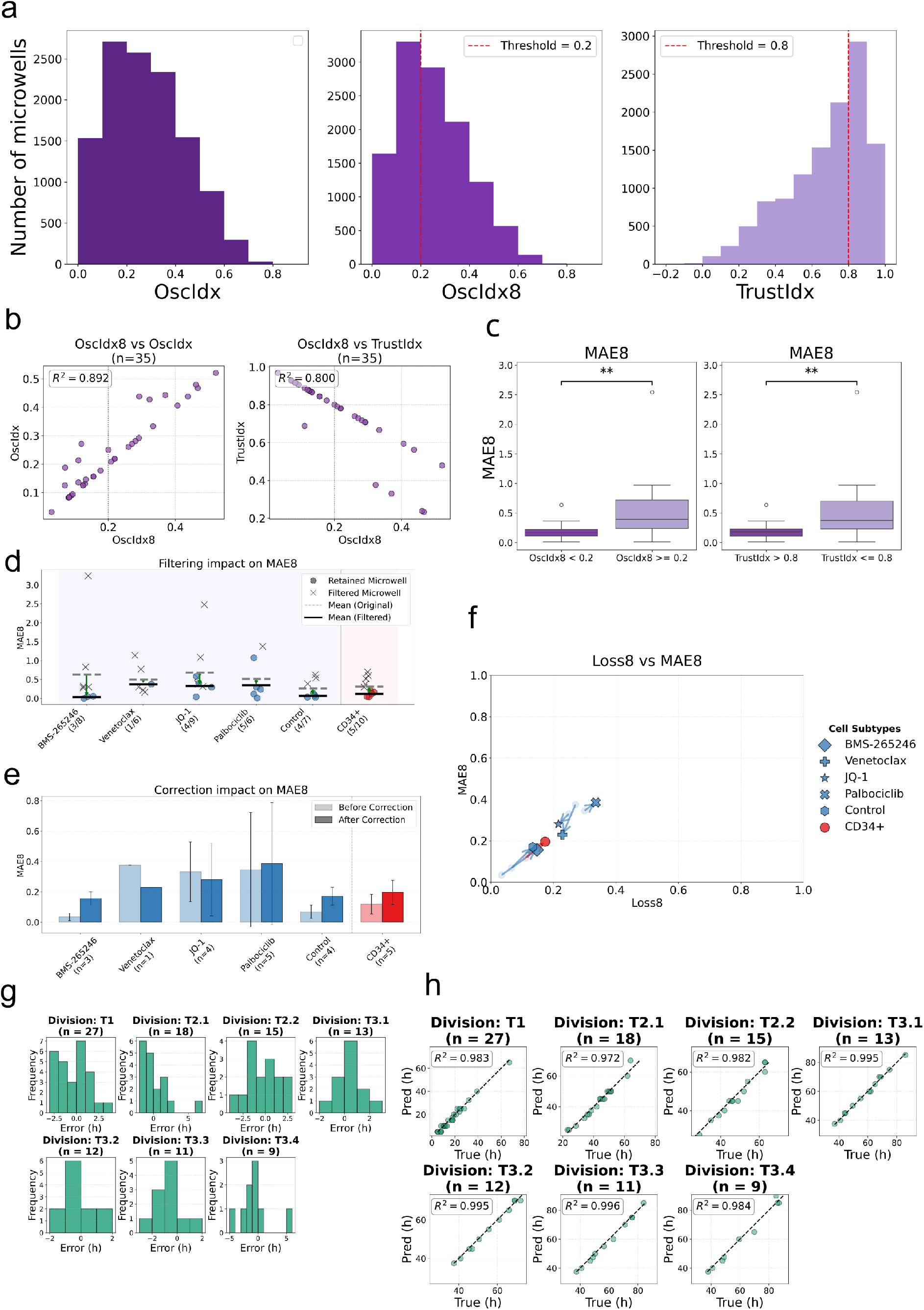
VisioWell quality control and comparison with manual annotated groundtruth **a)** Quality indexes distribution on complete dataset, merging OCI cell line and coord blood data and retaining only microwells for which at least one division was retrieved. **b)** Quality indexes, manually annotated groundtruth, n=35. A reference value of *OscIdx*8 = 0.2 is shown as a vertical gray line. **c)** MAE8 computed for microwells filtered with *OscIdx*8 *<* 0.2 or *TrustIdx >* 0.8. Mann-Whitney statistical test. **d)** Effect on *MAE*8 when filtering with quality index *OscIdx*8 = 0.2 for treated OCI cells (left) and primary cord blood cells (right). For each condition and type of cells, filtered microwells are represented with a cross, while retained microwells are represented with a full circle. The mean error for all microwells, before filtering, is represented with a dotted straight line. The mean error for the microwells after filtering is represented with a thick straight line. **e)** Effect of curve correction on *MAE*8, after filtering with *OscIdx*8 = 0.2. **f)** Scatter plot for OCI cell line and cord blood microwells of *Loss*8 and *MAE*8 before and after correction, results filtered with *OscIdx*8 = 0.2. Each symbol represents the average of the two coordinates across all microwells for that condition after correction. The circle represents the average of the two coordinates across all microwells for that condition before correction. An arrow connects the corresponding starting point, indicated by the circle, and the ending point, indicated by the symbol. **g)** Division time error distribution in hours, for all division times. Results filtered with *OscIdx*8 = 0.2 and corrected. **h)** Scatter plots of true vs predicted for all division times. Results filtered with *OscIdx*8 = 0.2 and corrected, using Pearson correlation.

We then evaluated the impact of filtering and curve correction on the segmentation, considering a larger, but still manually annotated dataset. This dataset included microwells containing more than one cell at T0, to quantify segmentation metrics regardless of the choices in the algorithm pipeline. Filtering microwells based on their *OscIdx*8 value (*<* 0.2) selected the highest quality wells for all tested conditions, as shown by the decreased *MAE*8 score. Nevertheless, conditions characterized by limited cell division and/or a higher incidence of cell death (Venetoclax, JQ-1 and Palbociclib), resulted in the selection of fewer microwells and/or a larger standard deviation of the *MAE*8 (Fig. 3d). When we introduced the change point curve correction, we observed a moderate oscillation of the *MAE*8 score in both directions, but always keeping the metric below 1 (1 hour), which we consider acceptable (Fig. 3e). Moreover, *MAE*8 normalized over the number of cells in the ground truth is smaller (Fig S2a). After filtering and correcting, both *MAE*8 and *Loss*8 (fraction of frames with incorrect cell count over the same frame window) values decreased below 0.2 (Fig. 3f). We also compared the finetuned CellposeSAM model with a finetuned YOLO model for object detection, and with zero-shot CellposeSAM and zero-shot microSAM, confirming better performance for finetuned CellposeSAM (Fig S2b-d). The choice of the statistical index used in curve correction did not introduce significant changes in MAE8 (Fig S2e). Additionally, CellposeSAM was evaluated for the whole duration of the acquisitions, with results similar to those obtained up to the third division (Fig S3a-c). Finally, we benchmarked the retrieval of division times: CellposeSAM retrieved division times displayed a moderate error distribution, comparable to the delay between each frame acquisition (Fig. 3g). When retaining only the clonal microwells (only one cell detected at T0), the *R*^2^ coefficient between predicted and true times showed a strong correlation, with values ranging between 0.971 and 0.996, (Fig. 3h). All these results support a high-quality automatic annotation process for VisioWell, able to analyse thousands of clones at once for their cell division length.

### VisioWell detects drug-induced cell-cycle perturbations in AML

After establishing the reliability of the image analysis pipeline, we tested if VisioWell could reliably measure the division dynamics of a biologically relevant model. We turned to human cord blood CD34+ HSPCs, which have been extensively studied for their cell division dynamics [25]. We tested the whole compartment, without subsetting into different HSPCs, and we observed an extended time to complete the first cell division, compared to the following divisions. This observation is consistent with previously reported behaviours for specific subsets, generalised for the entire compartment (Fig. 4a). Daughter cell division times (T2.1 vs T2.2) were highly correlated (Fig. 4b), in accordance with the known strong synchronicity observed across sister cells belonging to the same family [8]. These results confirm that VisioWell correctly retrieves the division dynamics in hundreds of healthy primary and motile HSPC-derived families, without the need for cell labeling.

**Figure 4:**
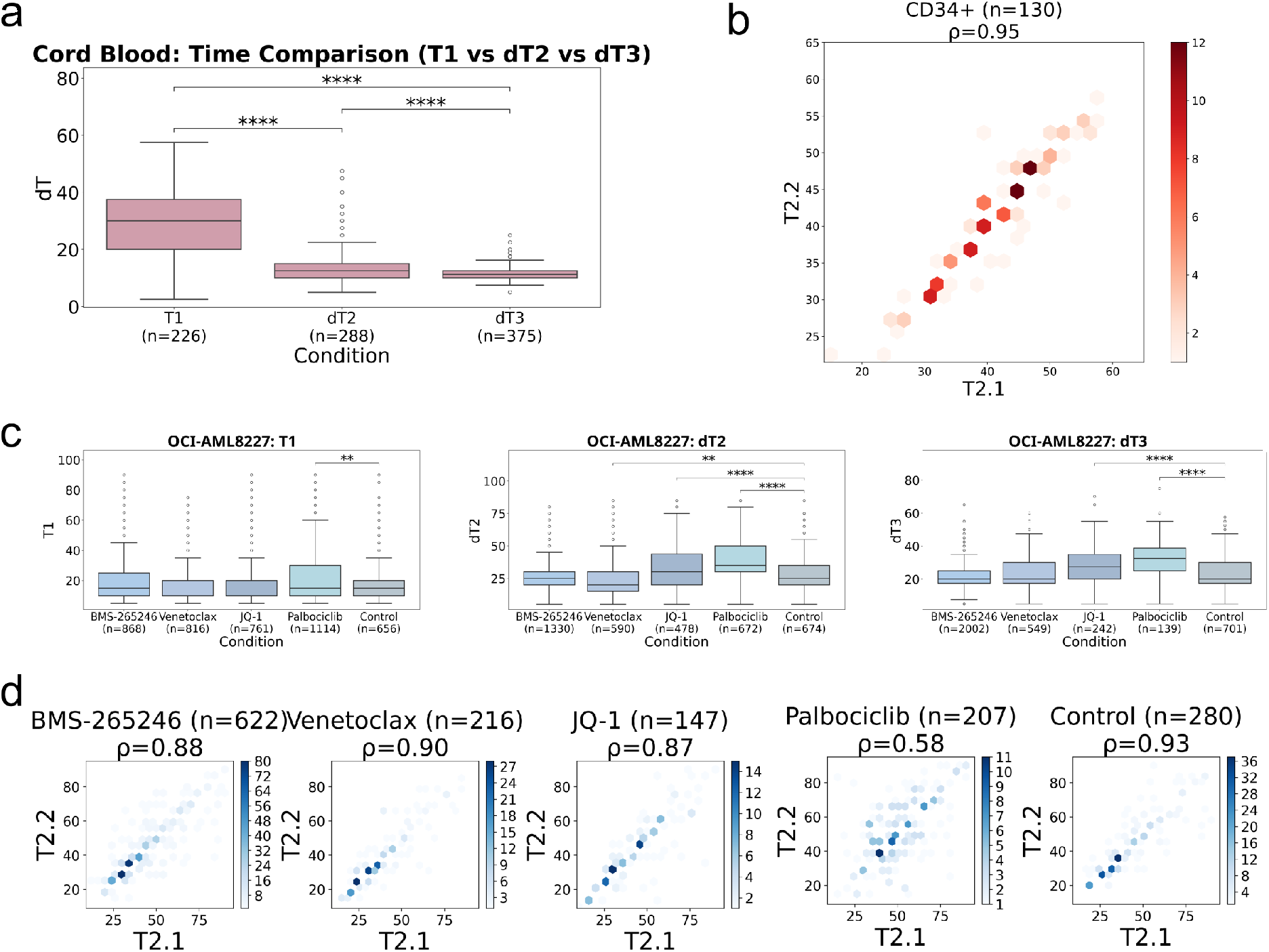
VisioWell experimental validation: **a)** Box plots for CD34+ cord blood cells for T1, dT2, dT3. Results filtered with *OscIdx*8 *<* 0.2 and *TrustIdx >* 0.8 and corrected. Statistical significance was assessed via the Kruskal-Wallis test and post-hoc Dunn test with Hochberg-Simes correction. **b)** Hexagon plot for CD34+ cord blood cells and Spearman correlation co-efficient between T2.1 and T2.2. Results filtered with *OscIdx*8 *<* 0.2 and *TrustIdx >* 0.8 and corrected. Each point represents a complete division of the mother cell into two daughter cells at times T2.1 and T2.2. **c)** Box plots for OCI-8227 T1, dT2, dT3. Results filtered with *OscIdx*8 *<* 0.2 and *TrustIdx >* 0.8 and corrected. Statistical significance was assessed via the Kruskal-Wallis test and post-hoc Dunn test with Hochberg-Simes correction. **d)** Hexagon plot for OCI cells and correlation between T2.1 and T2.2. Results filtered with *OscIdx*80.2 and *TrustIdx >* 0.8 and corrected. Each point represents a complete division of the mother cell into two daughter cells at times T2.1 and T2.2

Next, we investigated whether VisioWell could detect drug-induced perturbations in division dynamics. We treated the acute myeloid leukemia cell model OCI-AML8227 cells continuously, with four drugs that could potentially alter the cell-cycle length over 96 hours: BMS-265246 (CDK1/2 inhibitor)[26], Venetoclax (BCL2 inhibitor)[27], JQ-1 (BRD4-inhibitor)[8], and Palbociclib (CDK4/6 inhibitor)[25]. We detected an extended T1 only for the Palbociclib-treated cells, a result that was conserved across the entire range of divisions analysed (Fig. 4c). Unlike cord blood HSPCs, which displayed high sister-cell synchronicity (T2.1 vs T2.2), OCI cells displayed a significant loss of synchronicity specifically under Palbociclib administration (Fig. 4d). These results suggest a potential role for palbociclib in modulating cell division length for AML cells, and confirm the potential for VisioWell use in drug screening and validation.

## Discussion

Live-cell imaging to investigate haematopoietic cell biology is not a novel technique: reports from the early 2000s [2, 28–30] showcased the potential of this approach, highlighting properties that are still investigated twenty years later [8]. Those remarkable results required a high degree of technical expertise, both in microfabrication and image analysis, a rare occurrence in most experimental hematology labs. To overcome these limitations, we developed VisioWell to enable high-throughput end-to-end automated analysis of haematopoietic cell division dynamics. The array includes thousands of circular microwells, optimized to confine and trap non-adherent cells. The microwell design reduces the risk of external cells entering the microwell and prevents the cells already there from leaving. We used a thermoplastic elastomer that is robust, durable, biocompatible, and relatively inexpensive: to produce one hundred microarrays, we spent approximately one hundred euros to purchase the raw material. As an example, a 12-well plate array contains approximately 3350 microwells, and with the maximum loading efficiency estimated around 25-30%, it is possible to analyse more than 800 individual clones in a single well. Therefore, VisioWell offers an affordable and high-performance alternative to previous methods, enabling different experimental conditions in a single assay.

We demonstrated the VisioWell’s potential to analyse the first three cell divisions. Further development is required for analysing more divisions, as individual cell tracking becomes quite challenging computationally at the frame rate used for this study. The division retrieval logic relies on the assumption that sister cells complete their division before cousin cells start dividing, which may not hold true for different experimental settings. We encourage users to test this assumption beforehand. Another challenge is death detection, although the curve correction partially deals with dying cells that are intermittently detected by the segmentation network. We were careful to evaluate the quality of VisioWell output: we designed specific quality indexes, tuned on the specific dataset. They are able to discard poor-quality microwells and can be used as annotation-free filters. We also benchmarked the main steps of the computational pipeline (i.e., segmentation, curve correction, and time division retrieval) against manual annotation, with high correlation.

We tested VisioWell with healthy HSPCs, detecting a longer time to complete the first division and a high degree of synchronicity, as previously described [8], and overall validating our approach. We also tested the cell model OCI-AML8227, treated with sublethal doses of four molecules, with the goal of detecting any modification of cell division time and synchronicity. Only the CDK4/6 inhibition mediated by palbociclib affected both cell division properties. The extended time to divide across three divisions reflects the accumulation in early G1 [31], an expected consequence of G1/S checkpoint inhibition. The decrease in synchronicity was unexpected, though, and raises several exciting perspectives: it is established that a drug inducing an early G1 arrest, upon washout, leads to a general synchronisation of a cell population, as all those cells have been kept in the same cell cycle phase until the drug removal. Nevertheless, we could not find a study that looked at paired-daughter cell synchronicity in the presence of drug sublethal doses. One potential interpretation is that the drug sublethal dose only partially affects the G1/S checkpoint transition, suggesting that paired daughter cells can display different sensitivity to cell cycle inhibition. It would be interesting to investigate further if this differential sensitivity is a sign of the asymmetric distribution of the cellular machinery responsible for cell cycle control. On a different note, continuous CDK4/6 inhibition in healthy human HSPCs extends cell cycle length and can increase the percentage of CD34+ cells [25, 32, 33]. As OCI-AML8227 is a model for leukaemic stem cell (LSC) biology, and we also detect an extension in cell cycle length, it would be interesting to assess if CDK4/6 inhibition impacts the distribution of LSCs in a way that is similar to healthy HSPCs.

In conclusion, VisioWell is a novel experimental tool with screening potential, able to assess how drugs impact cell division synchronicity, providing a first crucial step into uncovering drug-induced cell division modulation. Moreover, as an all-in-one solution, it simplifies access to live-cell imaging experiments, adding a new, continuous dimension to single-cell functional assays.

## Experimental workflow

### Microwells array production

#### 1. Mould production

A starting brass mould should be produced via micromilling (design available upon request). While durable, routine use of the brass mould requires progressively longer cleaning steps with ethanol. We include here the procedure to produce a polyepoxy mould that is comparable in performance via PDMS negative, which could replace the brass mould for routine production.

i. In a Petri dish, cover the brass mould with 50g of silicone elastomer mix (1/10 of binding agent) and homogenise with a spatula.
ii. Place the mix in a vacuum bell until all the air bubbles are gone, then solidify the mix at 60 °C for at least 2 hours.
iii. Carefully unmould the PDMS from the mould using a scalpel to detach the edges without scratching the brass surface.
iv. Place this counter-mould in a vacuum bell for 1 hour, with 30 mL trichloro(perfluorooctyl) silane dispensed into 3 separate recipients. This step ensures homogeneous silane gas-phase deposition, which improves the durability of the negative mould.
v. Cover the lithography wafer with 50g of silicon elastomer + 1/10 binding agent mix and repeat the homogenisation, vacuum, and solidification steps for 1 hour.
vi. Cover the silanised counter-mould with 50g of epoxy resin (50% binding agent) and repeat the homogenisation, vacuum, and solidification steps for 1 hour.
vii. Unmould and separate the PDMS layer from the epoxy by slicing a thin line with a scalpel between both layers.

#### 2. Microwell array production and plate preparation

This step includes the creation of reproducible arrays via hot embossing and their incorporation within a tissue culture vessel. We provide a standardised procedure, which increases the entire process reproducibility, for a 12-well tissue culture plate, but it could be adapted to any other format, according to the specific measurements of each culture vessel.

i. Before starting, transfer the mould to an oven set at 145 °C.
ii. Cut the Flexdym in round patterns of 20 mm of diameter (or accordingly to the tissue vessel specifics). The use of a cutter plotter accelerates this step.
iii. From this step onwards, we suggest working in a dust-free environment (e.g., a PCR cabinet). Remove one side of the protective vinyl and stick the flexdym pattern onto a clean glass slide, while removing any air bubbles. Keep the protective vinyl aside in a clean Petri dish for further use.
iv. Remove the second protective vinyl, then transfer the Flexdym against the hot mould and secure the glass slide and the mould together using 2 metallic clips.
v. Leave the mould with the Flexdym for 30 seconds in the oven at 145 °C
vi. Take the chip out of the oven to the hood, quickly remove the metallic clips, and detach the mould. If some polymer remains on the mould, add a few drops of ethanol in between to help the unmoulding.
vii. To emboss another array, return the mould to the oven at 145 °C. If the embossing is finished, sonicate the mould in a 50/50 ethanol-acetone solution, then rinse with deionized water and store in a clean Petri dish.
viii. Under a microscope, check for complete embossing: a square network of microwells should be clearly visible, with no impurities in the middle and only on the edges of the array.
ix. Perform a plasma cleaning for 30 seconds on the glass slides with the embossed chip. This step should make the microarray surface hydrophilic and cell culture compatible.
x. Detach the microwell array from the glass slide using tweezers. Then, align the array with the tweezers and fix it at the bottom of the plate. Finally, take the vinyl sheet shed in step iii to cover the array, and press delicately but thoroughly on the array to ensure that no large air bubble is stuck between the tissue culture plate and the microarray.
xi. Leave the plate at 60 °C for 1 hour to ensure that the array is adhering to the bottom of the plate.
xii. Add 70% ethanol to the wells and place the vessel in a vacuum bell to remove any air bubbles left. From this step onwards, the plate should be handled in a sterile environment.
xiii. Wash each well with deionized water three times. It can be stored for weeks if adequately wrapped with plastic film.

### Cell seeding for live-cell imaging

We optimised the cell seeding concentrations for haematopoietic cells. For different motile cell types, we suggest testing a range of concentrations to maximise the number of microwells containing a single cell. As for the previous step, the provided volumes and concentrations are intended for a 12-well plate array. The cells of interest can be FACS-isolated before seeding, or directly FACS-sorted onto the microwell array (see “alternative seeding”).

i. Remove the deionized water and rapidly rinse with 1 mL of PBS 1X. Repeat this step two more times, carefully ensuring that the microwells are never completely dry, as this could lead to the formation of air bubbles.
ii. Replace the PBS 1X with 900 µL of the final cell culture medium.
iii. Resuspend between 5×103 and 1×104 cells of interest in 100 µL of cell culture medium.
iv. Carefully add the cells dropwise, making sure to add each drop in a different part of the well surface. Then, gently but rapidly agitate the plate horizontally (once) and vertically (once), to ensure homogeneous distribution.
v. We strongly encourage to perform the cell seeding close to the imaging device, and load the plate immediately after seeding for acquisition. This should improve the settling of the seeded cells in the microwells, as transporting the plate could alter the cells’ sedimentation, leading to suboptimal seeding

#### Alternative seeding

it is also possible to directly FACS-sort the cells of interest onto the microwells array. While this approach could reduce cell loss for a rare individual population, there are several drawbacks: the cell culture medium could be mixed with the sheath buffer used by the cell sorter, especially for high cell numbers, and the cell sedimentation steps are less standardised and susceptible to experimental variability.

### Live single cell imaging and data retrieval

While VisioWell can work with any microscope or imaging device adapted to live-cell imaging, we strongly suggest using a platform that allows manual setting of the field focus. This is to avoid any focus loss due to the microwell array, which may be imperfectly fixed to the bottom of the tissue culture plate well. We suggest waiting between 1 and 2 hours before starting the acquisition to ensure cell sedimentation.

i. Carefully load the tissue culture vessel with the microwells array.
ii. Select the brightfield imaging option, setting a 10 ms exposure time. Verify the quality of the images actively and adapt the light intensity accordingly.
iii. Set the focus on the bottom of the array, making sure to select areas that contain microwells with at least one cell.
iv. Set the image acquisition frequency and length according to the experimental needs. While for the 12-well format the evaporation risk is minimal, we do not exclude that a longer timescale can affect the cell culture parameters for smaller volumes. We suggest avoiding disturbing the plate during the acquisition to prevent any cell resuspension.
v. Once concluded, the data can be exported as a single .czi file.

### Data analysis

1. **Running the time division retrieval algorithm** Note: the execution of the algorithm requires the prior installation of Python and the required packages (GitHub link: https://github.com/tizianatocci/VisioWell).
  i. Step 0: microwells identification: activate the Python environment for identification of microwells (envA). Launch the code step0 findwells.py from the terminal, indicating the folder containing the video to analyse, the rotation angle, and the folder in which to save the results.
  ii. Step 1: cell segmentation and curve correction: activate the Python environment for cell identification and time division retrieval (envB). Launch the code step1 predictions.py from the terminal, indicating the output folder of step 0, the folder in which to save the results, and the name of the training to load.
  iii. Step 2: time division retrieval: activate the Python environment for cell identification and time division retrieval (envB). Launch the code step2 output.py from the terminal, indicating the output folder of step 1, the folder in which to save the results, the cell type, the imaging frequency, the output folder of step 0, and the path in which to save QC report.
  iv. Visualization of results: Activate the Python environment for visualization of results (envC). Run the notebook results.ipynb
  v. Eventual retraining of neural network with custom images: Obtain manually the masks of all the cells in your frame. Run the code training.py.

## Methods

### Microwell array

The custom microarray consists of an array of thousands of microwells (1225 microwells per cm²). Each microwell is 100 µm large and 150 µm deep. The upper 50 µm of the microwell is arc-shaped. The microwell design is micromilled on a brass plate for optimal precision and durability, and could be replicated on epoxy. Microarrays are produced by hot embossing the mould onto pre-cut Flexdym sheets. The array sizes are compatible with 6-, 12-, and 24-well tissue culture plates. After embossing, a plasma treatment is applied to the array, as the resulting hydrophilic surface ensures the homogeneous distribution of the culture medium. The arrays are then fixed to the bottom of the plate wells, preventing air bubble formation and uneven surfaces. The assembled plates are then sterilised, rinsed, and stored in sterile water (Fig. 2b). This procedure can be completed in less than 2 hours, and plates can be stored for several weeks at + 4 °C.

### Microwells design and moulding

The microwells mould was designed using Fusion 360 (Autodesk) and manufactured from brass using a CNC micromilling system (Optimum Optimill MH 22Vd). The template is available upon request. The resulting brass mould was used as the template for subsequent replica moulding. A negative mould was prepared using polydimethylsiloxane (PDMS; Sylgard 184 Silicone Elastomer Kit) base and curing agent mixed at a 10:1 ratio, poured onto the brass mould, degassed for 15 min under vacuum, and cured at 60 °C for 3 h. After curing, the PDMS negative replica was carefully peeled off, then silanized in a vacuum chamber using 10 trichloro(perfluorooctyl)silane (Sigma-Aldrich) for 1 h. Alternatively, an epoxy positive mould was generated by casting epoxy resin following the manufacturer’s instructions (EpoxAcast 670 HT, Smooth-On), applying uniformly 20g of epoxy resin with curing agent into the PDMS cast with a soft brush, degassing, and curing for 3 h.

### Flexdym patterning

We used Flexdym, a flexible polymer sheet (Edentech), to produce the microwell chips. 250 µm-thick sheets were cut into circular formats adapted to 24-, 12-, or 6-well plates using a cutter plotter (Graphtec system) with predefined vector designs (Adobe Illustrator). Flexdym sheets were laminated onto clean glass slides (Superfrost, Thermo Fisher Scientific) and secured with 25mm metallic clips and heated at 145 °C for 5 to 30 seconds, depending on the chip format. After the hot embossing, chips were detached under a laminar flow hood, using ethanol to facilitate the release. Embossed chips were inspected under a microscope to confirm proper pattern transfer before plasma cleaning for approximately 30 seconds (Harrick Plasma). Chips were then carefully aligned to the bottom of the culture plate by pressing them down and incubating for 1 h at 60°C. Plates were then sterilised by filling the wells with 70% ethanol under a vacuum bell for 15 minutes, then rinsed 3 times with sterile ultrapure water and stored at 4 °C in sterile water for further use.

### Ethical approval and processing of human samples

De-identified umbilical cord blood units were collected according to the guidelines defined by the Saint-Louis Hospital Umbilical Cord Blood biobank (authorization AC-2016-2759) and in accordance with the Declaration of Helsinki. Mononuclear Cells (MNC) were isolated via Ficoll density gradient, then underwent CD34-enrichment using immunomagnetic microbeads (MACS Miltenyi). CD34-enriched fractions were immediately processed or frozen in Fetal Bovine Serum (FBS, Eurobio Scientific) supplemented with 10% dimethyl sulfoxide (DMSO, Sigma) for further analysis.

### Fluorescence Activated Cell-Sorting, cell culture and drug treatment in microwells

Enriched CD34+ fractions, both from cord blood or OCI-AML8227, were stained with the antibody CD34 APC-Cy7, clone 581 (BioLegend) added at a 1:40 dilution. All samples were incubated for 40 min at 4°C, then washed and resuspended in PBS. Live/dead discrimination was performed with DAPI staining at a final concentration of 1 µg/mL. Cell populations were sorted using a BD Fusion cell sorter (Becton Dickson Biosciences), and data analysis was performed with FlowJo v10 (Waters Biosciences). The OCI-AML8227 cells were maintained in StemSpan II (StemCells) supplemented with Pen/Strep 1x (Gibco), SCF at 50 ng/mL (Peprotech), FLT3L at 50 ng/mL (Peprotech), TPO at 25 ng/mL (Peprotech), IL-6, IL-3 and G-CSF at 10 ng/mL (Peprotech) and passaged twice a week at 3 × 10^5^ cells/mL. Cells were tested negative for Mycoplasma contamination using MycoStripTM 50 (InvivoGen, rep-mysnc-50). Primary HSPCs were cultured with the following medium: StemPro-34 SFM (ThermoFisher Scientific) supplemented with Penicillin – Streptomycin 1X (ThermoFisher Scientific), L-Glutamine 2 mM (ThermoFisher Scientific), SCF 100 ng/mL, TPO 100 ng/mL, IL-6 50 ng/mL (Peprotech), GM-CSF (Miltenyi) 25 ng/mL, FLT3l 20 ng/mL, EPO 1 U/mL (Janssen) and IL-3 10 ng/mL. For the drug screening experiments, compounds such as venetoclax (Sigma) at 5 nM, JQ1 (Sigma) at 5 nM, palbociclib (Sigma) at 50 nM, and BMS-265246 (Sigma) at 500 nM were added at sublethal doses, defined as the lowest dose inducing a cell viability of at least 90%.

### Live-cell imaging

Plates were loaded into a Cell Discoverer 7 (Zeiss), with the imaging chamber set at 37 °C, 5% CO2. Acquisitions were conducted with 30 mn intervals between frames, for a total of 96 hours of time-lapse imaging. Brightfield acquisition was conducted with 50% illumination intensity and 10 ms exposure time using Zen software. Multiple tile regions were defined along the x and y axes to cover the microwell arrays, and focus positions were calibrated manually across the z-axis. Three independent experiments were conducted: two replicates for the OCI cell line (frame rate: 1 hour over 96 hours) and one experiment for cord blood cells (frame rate: 30 minutes over 61 hours).

### Computational requirements

The computational method has been implemented in Python 3.10, leveraging the following main libraries: Cellpose, PyTorch, cupy, numpy, scipy, OpenCV, ruptures. GPU acceleration is leveraged for extensive matrix computations for microwell identification and for segmentation.

### Image preprocessing

The slide acquisition video eventually undergoes a rotation to align the microwells with the x-axis to correct for significant angular tilts. A drift correction algorithm based on cross-correlation is employed to improve the registration between frames. Then, each frame is converted to 8 bit appropriately adjusting the pixel range to improve the contrast.

### Well identification and labelling

The Circular Hough Transform (CHT) algorithm [34] is used to detect the circular microwells. By changing the parameters concerning the radius of the well to be identified, the algorithm can be applied to frame acquisitions with different magnification and possibly different well sizes across experiments. As the microwells are organized in columns and rows in the microfluidic chip, in each field of view the naming convention consists of identifying rows with letters in alphabetical order and assigning numbers to each column. Once the centers of the microwells are identified, all the possible lines passing through each pair of points are found. The angles between the x-axis and the identified lines are computed, and only the lines with an angle under a threshold are retained. Groups of parallel lines are retrieved and only the group of lines with the highest number of lines and smaller angle is selected. Groups of overlapping lines sharing at least one point are found, and inside each of these groups only the longest line is kept. The points belonging to the lines are found based on the distance of each point from the line. Finally, the points are sorted inside each line based on their x, y coordinates, and a name composed of a letter and a number is assigned to each microwell.

### Neural networks training for cell segmentation

The cells are segmented with a fine-tuned CellposeSAM model. The segmentation algorithm is initially run to detect the number of cells in the first frame. If no cells or more than one cell is detected, the microwell is discarded. For training, the cell masks were manually annotated with the Cellpose GUI, excluding dead cells, i.e., cells that lose their morphological integrity and cease to move. The total number of manually annotated frames is 1209, including both OCI-8227 and cord blood cells. Each frame contains a variable number of cells, ranging from isolated single mother cells to saturated microwells. A hyperparameter search was conducted with the hyperopt library [35] to select the best learning rate and weight decay hyperparameters, and MLFlow [36] was used for reproducibility and tracking of training. CellposeSAM pretrained models were fine-tuned for 50 epochs with a batch size of 8 on images of size 256*px ×* 256*px*. The built-in data augmentation was leveraged.

### Alternative approach for cell identification: neural network for object detection

While the core of the pipeline is based on segmentation and specifically on CellPoseSAM, an alternative version of the algorithm based on Object Detection, leveraging the YOLO v11 ultralytics implementation [37], was developed. All the other steps, namely the well identification step, the curve correction, and time retrieval, remained unchanged. The segmentation core was replaced with an object detection core. An algorithm was used to automatically obtain the bounding box for the YOLO training from the manually annotated cells. Similarly to CellPoseSAM training, a hyperparameter search was conducted with the hyperopt library to select the best learning rate and weight decay hyperparameters, and MLFLOW was used for reproducibility and tracking of training. YOLO v11 pretrained models were fine-tuned for 50 epochs with a batch size of 8 images of size 256*px ×* 256*px*. The built-in data augmentation was leveraged.

### Alternative approach for cell segmentation: zero-shot CellposeSAM and microsam

For additional comparison, predictions were obtained with CellposeSAM [22] and microsam zero-shot [38], i.e., no additional fine-tuning was performed.

### Curve correction

The Python package “ruptures” for change point detection in time series data is employed to implement the curve correction post-processing method [23]. The “pelt” algorithm with a penalty of 2 is used to find the optimal segmentation of the time series, i.e., the cell count output, by minimizing the L2 norm within each segment. Once the segments are identified, the oscillations are flattened based on the segment’s most likely value, MLV. The MLV for each segment *n* is computed as follows:

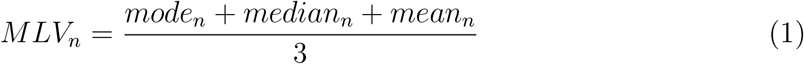

where *mode_n_*, *median_n_* and *mean_n_* are respectively the mode, median and mean of the cell count output for the segment *n*. Only microwells with one cell after curve correction are retained to automatically retrieve division times for the downstream analysis. In (Fig S2e), the curve correction was implemented relying only on the mode, median, and mean. For each case, the difference between the MAE8 computed with a single statistical index and the MLV was computed.

### Division times retrieval

The first seven times of divisions are automatically retrieved, and they are indicated as T1, T2.1, T2.2, T3.1, T3.2, T3.3, T3.4. The division times are automatically computed from the corrected cell counting output, which contains the numbers of cells in each frame after correction. When the number of divisions increases, a division time is automatically identified. For the first increase (one to two cells), the division time T1 is assigned by multiplying the number of frames for which the increase is observed by the time interval between frames. If the increase is bigger than one cell (e.g. two sister cells dividing into four cousin cells), then multiple simultaneous and overlapping times of division are assigned (in this case, T2.1 and T2.2 are co-occurring). Cell death is detected when the cell corrected count decreases; it is appropriately handled by setting to NA the times of division corresponding to the dying cell and to its hypothetical future descendants. For example, if T2.2 is set NA due to a daughter cell dying, then T3.3 and T3.4 are set to NA as well.

### Quality Index and Filtering

The quality index serves as a metric to assess the level of oscillations in the cell detection output. The oscillation index is defined as follows:

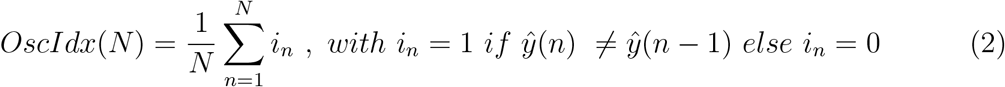

where *ŷ* is the predicted number of cells. *OscIdx* is defined for *N* representing the total number of frames, while *OscIdx*8 is defined for *N* denoting the last frame for which the corrected output indicates the presence of eight cells. *OscIdx*(*N*) is a measure between 0 and 1 and it computes how many oscillations occur in the cell detection output before correction. The results are filtered according to the *OscIdx*8 in order to keep only the most reliable cell quantifications.

### Quality Report

The quality report delivers a quantification of the algorithm performance for one whole experiment according to the metrics defined. The metrics are reported for both microwell identification and time retrieval. The quality control for microwell identification includes: number of scenes retrieved, total number of microwells found, mean number of microwells found per frame, predicted percentage of loading efficiency, list of the frames having a naming out of range [A-E; 1-5]. The quality control for time retrieval includes: number of microwells analyzed, percentage of microwells with a *TrustIdx <* 0.5, percentage of microwells with *OscIdx*8 *> threshold*, percentage of microwells with *Colony Size >* 30 (CSL, which may be due to intruder cells), and percentage of microwells with generation discrepancies (GD), i.e., not enough time passed between the first division of a cell generation and the last division of the previous generation. The thresholds for CSL and GD can be set by the user according to the specific experimental setting. The metrics are defined as follows:

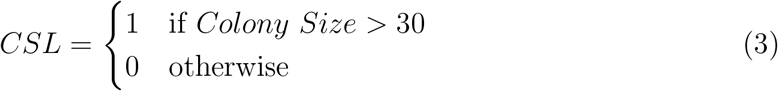

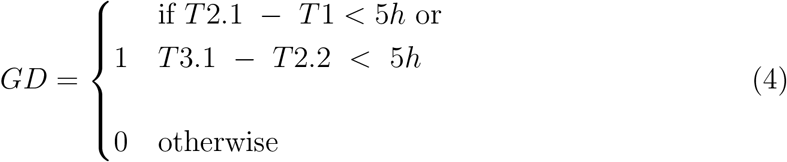

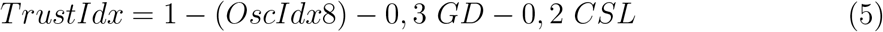

A Streamlit dashboard included in the algorithm enables to easily verify the results obtained. The section for microwell identifications allows the user to verify the naming of frames by inserting the folder and file names. The section for segmentation results loads the video of the entire well and it overimposes the segmented masks. Finally, the section for time division retrieval loads the microwell frames and the cell number-frame curve around the eventual generation discrepancy.

#### Performance metrics

For performance quantification, the metrics Mean Absolute Error (MAE) and Loss are defined as follows:

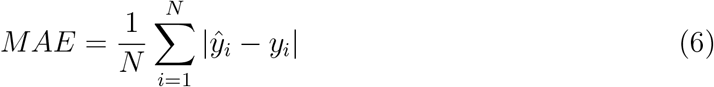

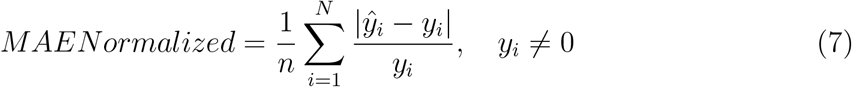

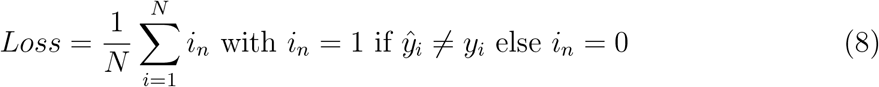

where *ŷ_i_* is the predicted number of cells, *y_i_*is the groundtruth number of cells in frame *i* and *N* is the number of frames.

#### Analysis on quality indexes

The dataset used for representing quality indexes (Fig 2h) includes 35 samples across OCI-8227 (28) and cord blood microwells (7). OCI cell line samples include 6 BMS microwells, 4 Venetoclax microwells, 8 JQ-1 microwells, 3 Palbo microwells, 7 Control microwells. Cord blood samples include 7 CD34+ (Hematopoietic Progenitor Cell) microwells.

The MAE8 was computed as specified in the previous paragraph Performance metrics. A microwell was labelled unreliable if *MAE >* 0.3 or if *MAE >* 0.4. Independently from the chosen cutoff (*OscIdx <* 0.2 and *TrustIdx >* 0.8), the two indexes *OscIdx* and *TrustIdx* were used to compute ROC-PR curves against the unreliability label. At the chosen thresholds (*OscIdx <* 0.2 and *TrustIdx >* 0.8), the recall (fraction of unreliable microwells identified) and the false positive rate (fraction of reliable microwell discarded) were computed. As a sensitivity analysis, the thresholds were re-selected by repeated stratified 5-fold cross-validation (200 repeats) on the 35 microwells: in each fold, the threshold maximizing Youden index (recall minus false-positive rate) on the four folds microwells was applied to the remaning held-out fold. The median selected thresholds across repeats were computed.

#### Groundtruth datasets

The groundtruth dataset used for segmentation benchmark (Fig 3a-c), before and/or after correction, was constructed by manually retrieving cell number of each frame from the live-cell acquisitions. To build the segmentation dataset, microwells are retained even if the algorithm identifies less or more than 1 cell in the first frame, to drop the strict requirement leveraged in the workflow (i.e. the first frame must contain one cell) and evaluate the segmentation capabilities of the system regardless of the downstream analysis. The segmentation benchmark dataset includes 46 samples across OCI (36) and cord blood microwells (10). OCI cell line samples include 8 BMS microwells, 6 Venetoclax microwells, 9 JQ-1 microwells, 6 Palbo microwells, 7 Control microwells. Cord blood samples include 10 CD34+ microwells.

The groundtruth dataset used for division times benchmark (Fig 3d-e) was constructed by manually retrieving division times from the live-cell acquisitions. The division times benchmark dataset includes 27 samples across OCI (22) and cord blood microwells (5). OCI cell line samples include 4 BMS microwells, 4 Venetoclax microwells, 3 JQ-1 microwells, 6 Palbo microwells, 5 Control microwells. Cord blood samples include 5 CD34+ microwells.

For obtaining the results of the comparison between multiple models, (Fig S3), the same microwells are considered, with no filtering.

#### Computation dT

A delay dT2 is defined by subtracting T1 from the division times of the daughter cells (*dT* 2.1 = *T* 2.1 *− T* 1; *dT* 2.2 = *T* 2.2 *− T* 1). A delay dT3 is defined by subtracting the average of the daughter cells division times (*avg_dT* 2) from the division times of the cousin cells (*dT* 3.1 = *T* 3.1 *− avg_T* 2; *dT* 3.2 = *T* 3.2 *− avg_T* 2,*dT* 3.3 = *T* 3.3 *− avg_T* 2,*dT* 3.4 = *T* 3.4 *− avg_T* 2).

### Statistical tests

The Pearson correlation test was used to compute the *R*^2^ values between quality indexes values and for the true and predicted division times. The Spearman rank correlation coefficient was computed between T2.1 and T2.2, after randomizing the order of observations. The Mann Whitney test was used to assess significant difference of MAE after thresholding based on *OscIdx*8 and *TrustIdx*. Statistical tests among OCI-8227 cell experimental conditions were performed with the Kruskal Wallis test and posthoc Dunn test. For multiple comparisons, the Hochberg-Simes correction is adopted.

### Data and code availability

Pre-processed live cell imaging datasets for reanalysis and reproducibility effort are available upon request to the lead author, as their size require appropriate and adequate strategies for long-term storage and sharing. The entire analytical pipeline is available on GitHub, together with the processed datasets: https://github.com/tizianatocci/VisioWell. The microwells mould template is available upon request to the corresponding author.

## Acknowledgements

We are indebted to Dr. Jan Audric and the IPGG technological core facility for their expertise in microfluidic device fabrication and their technical support. Equally, we would like to thank Dr. Louise Bonnemay and Dr. Benôıt Sorre for their assistance and for teaching us the intricacies of microfluidics. We would like to acknowledge Prof. Cristina Lo Celso and Dr. Constandina Pospori for sharing the OCI-AML8227 cells with us. We are grateful to the members of the Team Períe for their discussion, support, and their contribution to the manual annotation steps, particularly Margot Le Bot. We would like to thank the members of the Institut Curie Flow Facility for their help with setting up the flow cytometry experiments. We thank the members of the Institut Curie Imaging Platform for their assistance with the live-cell imaging experiments. We thank the Saint Louis Hospital Cord Blood Biobank for providing the biological resources necessary for this project.

## Author contributions

M.B. produced the microwell arrays and performed the live-cell imaging experiments. T.T. designed the analytical framework with the contribution of M.B. and performed the data analysis. E.M. and A.M. advised on the design of the analytical framework. H.I., L.P. and A.D. conceived the technological platform and provided overall supervision for the project. T.T., A.D. and M.B. wrote the manuscript, with substantial contribution from L.P. All the coauthors reviewed the manuscript and provided feedback.

## Supplementary figures

**Figure Supplementary 1:**
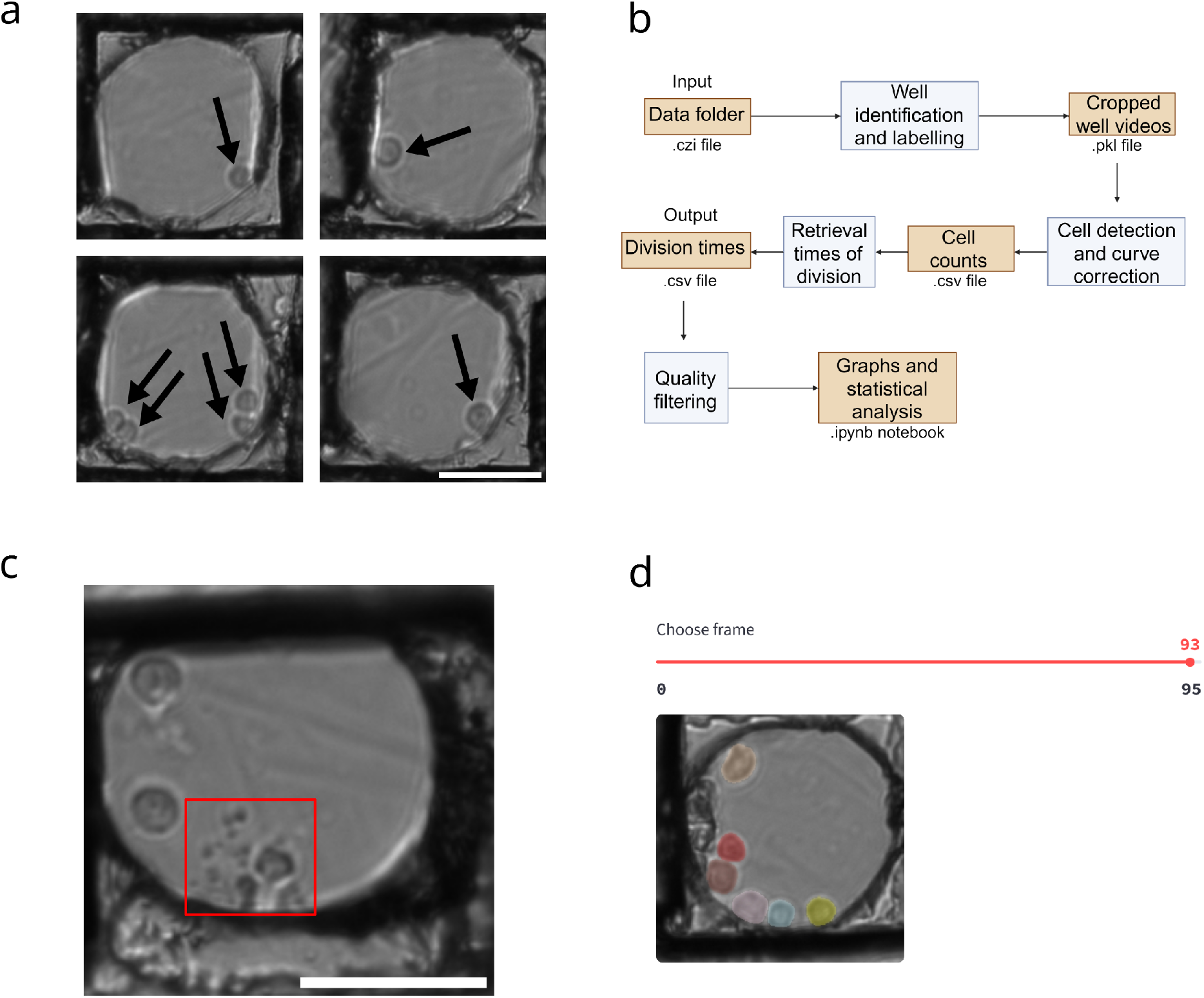
VisioWell platform. **a)** Cells sedimented in four microwells at T0 (beginning of the experiment). Scalebar: 50 *µm*. **b)** Flow diagram, describing the files generated at each step. **c)** Example of a microwell with a dying cell, indicated with a red rectangle. Scalebar: 50 *µm*. **d)** Streamlit app to visualize results. The user can choose the microwell to display and each segmented cell is indicated with a different color.

**Figure Supplementary 2:**
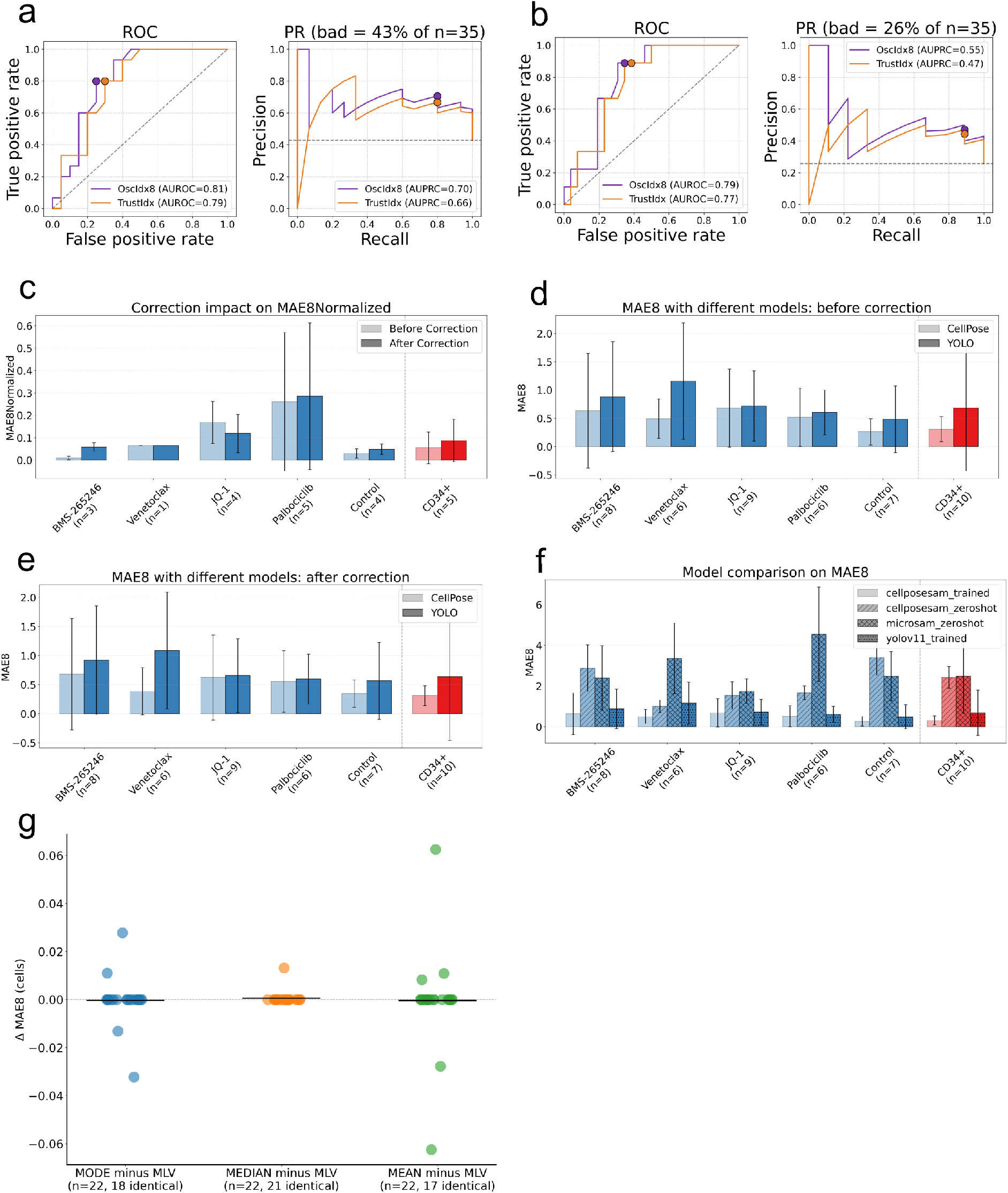
Additional pipeline validation. **a)** ROC and PR curves of *OscIdx*8 and *TrustIdx* for detecting unreliable microwells (*MAE >* 0.3). **b)** ROC and PR curves of *OscIdx*8 and *TrustIdx* for detecting unreliable microwells (*MAE >* 0.4). **c)** MAE8 normalized computed after correction. Histograms represent the *MAE*8 normalized, for each condition. **d)** Comparison for cell identification between CellposeSAM and YOLO before correction. Histograms represent the *MAE*8 for each condition. **e)** Comparison for cell identification between CellposeSAM and YOLO after correction. Histograms represent the *MAE*8 for each condition. **f)** Comparison for cell identification between trained CellposeSAM, zero-shot CellposeSAM, zero-shot microsam and trained YOLO V11. No correction applied. Histograms represent the *MAE*8 for each condition.

**Figure Supplementary 3:**
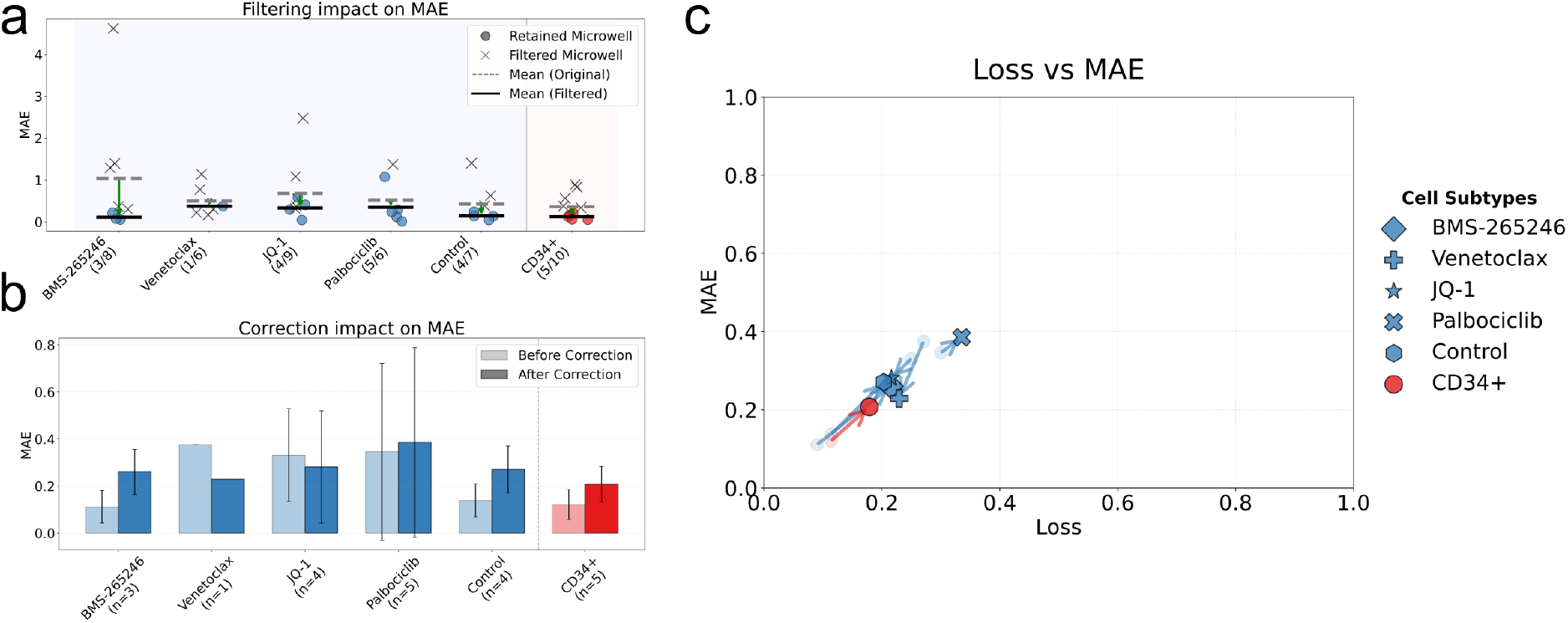
Segmentation benchmark for cells retrieved in all frames with the CellposeSAM algorithm. **a)** Effect on *MAE* when filtering with quality index *OscIdx*8 = 0.2 for OCI cells (left) and primary cord blood cells (right). For each condition and type of cells, filtered microwells are represented with a cross, while retained microwells are represented with a full circle. The mean error for all microwells, before filtering, is represented with a dotted straight line. The mean error for the microwells surviving filtering is represented with a thick straight line. **b)** Effect of curve correction on *MAE* on results filtered with *OscIdx*8 = 0.2. **c)** Scatter plot for OCI cell line and cord blood microwells of *Loss* and *MAE* before and after correction, results filtered with *OscIdx*8 = 0.2. Each symbol represents the average of the two coordinates across all microwells for that condition after correction. The circle represents the average of the two coordinates across all microwells for that condition before correction. An arrow connects the corresponding starting point, indicated by the circle, and the ending point, indicated by the symbol.

